# Benchmarking AlphaMissense Pathogenicity Predictions Against Cystic Fibrosis Variants

**DOI:** 10.1101/2023.10.05.561147

**Authors:** Eli Fritz McDonald, Kathryn E. Oliver, Jonathan P. Schlebach, Jens Meiler, Lars Plate

## Abstract

Variants in the cystic fibrosis transmembrane conductance regulator gene (*CFTR*) result in cystic fibrosis – a lethal autosomal recessive disorder. Missense variants that alter a single amino acid in the CFTR protein are among the most common cystic fibrosis variants, yet tools for accurately predicting molecular consequences of missense variants have been limited to date. AlphaMissense (AM) is a new technology that predicts the pathogenicity of missense variants based on dual learned protein structure and evolutionary features. Here, we evaluated the ability of AM to predict the pathogenicity of CFTR missense variants. AM predicted a high pathogenicity for CFTR residues overall, resulting in a high false positive rate and fair classification performance on CF variants from the CFTR2.org database. AM pathogenicity score correlated modestly with pathogenicity metrics from persons with CF including sweat chloride level, pancreatic insufficiency rate, and *Pseudomonas aeruginosa* infection rate. Correlation was also modest with CFTR trafficking and folding competency *in vitro*. By contrast, the AM score correlated well with CFTR channel function *in vitro* – demonstrating the dual structure and evolutionary training approach learns important functional information despite lacking such data during training. Different performance across metrics indicated AM may determine if polymorphisms in CFTR are recessive CF variants yet cannot differentiate mechanistic effects or the nature of pathophysiology. Finally, AM predictions offered limited utility to inform on the pharmacological response of CF variants i.e., *theratype*. Development of new approaches to differentiate the biochemical and pharmacological properties of CFTR variants is therefore still needed to refine the targeting of emerging precision CF therapeutics.

## Introduction

Cystic fibrosis (CF) is a lethal genetic disease caused by variants in the epithelial anion channel cystic fibrosis transmembrane conductance regulator (CFTR)^1^. CFTR is composed of an N-terminal lasso motif, two nucleotide binding domains (NBDs), two transmembrane domains (TMDs) and an unstructured regulatory domain (RD)^2^. Loss of CFTR protein production or function results in osmotic dysregulation at the epithelium of the skin, pancreatic duct, and lungs – leading to high sweat chloride levels, pancreatic insufficiency, and lung infections respectively^3^. Standard treatment paradigms for CF involve supplementation of salt, vitamins, and digestive enzymes, together with airway clearance therapies and small molecule CFTR modulators known as *potentiators* and *correctors*. CFTR variants experience distinct structural defects and proteostasis states leading to divergent pharmacological response profiles to modulators also known as *theratypes*^4–7^.

At present, elexacaftor-tezacaftor-ivacaftor (ETI) is the best available highly effective modulator therapy for CF. This triple combination is clinically approved for ∼170 CFTR variants, including the most commonly reported allele, deletion of phenylalanine 508 (F508del)^8–11^. ETI is composed of one gating potentiator (ivacaftor, VX-770) and two protein maturation correctors, tezacaftor (VX-661) and elexacaftor (VX-445). The corrector compounds have been suggested to directly bind unique subdomains of CFTR: VX-661 to TMD1^12,13^, and VX-445 to the N-terminal lasso and TMD2^14,15^. Correctors contribute intermolecular interactions that favor the properly folded, trafficking competent state of CFTR. Due to the distinct binding sites, VX-661 and VX-445 elicit different mechanisms of action and confer variable theratype responses. Thus, profiling CFTR variant theratypes to these and other emerging modulators remains an important priority for CF personalized medicine.

Increasing implementation of next-generation sequencing approaches for CFTR DNA analysis has rapidly augmented the pace of novel CFTR variant discovery; and thus, hastened the need for more accurate pathogenicity prediction tools. This is particularly relevant to individuals with CFTR related metabolic syndrome (CRMS), also known as CF Screen Positive Inconclusive Diagnosis (CFSPID). Patients are diagnosed with this condition if they possess a positive newborn screen for CF and either of the following criteria: (1) normal sweat chloride value (<30 mEq/L) and two identified CFTR variants, at least one of which exhibits unclear phenotypic consequences; or (2) intermediate sweat chloride value (30-59 mEq/L) and detection of one or zero CF-causing variants^16^. Clinical symptoms worsen for approximately 11-48% of CRMS/CFSPID patients, who eventually convert to a CF diagnosis^17,18^. Insufficient data exists to predict which CFTR variants (or other factors) enhance the risk for progression to CF.

Furthermore, high-throughput methods for characterizing CFTR variant severity are limited. Only 804 of the reported 2,111 variants have been annotated for disease association according to *in vitro* or clinical data^19^. The majority of these CFTR variants are single amino acid substitutions or missense variants^20^. Recently, AlphaMissense (AM) was published as a technology designed to predict the pathogenicity of missense variants throughout the human proteome^21^. Among well-characterized genetic diseases, AM included CF pathogenicity predictions for every possible CFTR single amino acid substitution. AM provides a significant advance beyond previous attempts to model a limited number of CFTR variants^22,23^. Here, we evaluated the predictive validity of AM across several metrics of CF data such as pathogenicity in people with CF, *in vitro* CFTR folding and function, and theratype.

The increasing pace of novel CFTR variant discovery has created a need for pathogenicity prediction, especially among wild-type (WT) heterozygous individuals, e.g. carriers, and CRMS/CFSPID individuals whose variants remain uncharacterized. Our analysis suggests AM predicts the relative pathogenicity of severe CF-causing variants well, while performing modestly for variants of unknown significance (VUS) or variants of varying clinical consequence (VVCC). Overall, AM showed a high false positive rate for predicting CFTR2 patient outcomes^19^. Among VUSs, two variants from CFTR2 and 368 variants from ClinVar^24^ were predicted as pathogenic. By contrast, the S912L VUS from CFTR2 was predicted benign despite clinical outcomes indicating ∼half the people with this variant display hallmarks of CF disease. AM scores correlated modestly with CF pathogenicity metrics and CFTR trafficking/folding competency *in vitro*. Correlation improved when compared to CFTR channel functional data. These analyses imply AM has learned important trends in variant function despite not training on such data. Finally, we provide evidence that AM offers little power in predicting CFTR variant theratype, although we note this measure is beyond its intended design. Thus, AM may offer capabilities in predicting the pathogenicity of emerging variants but proved less useful for theratyping variants.

## Results and Discussion

### I. AlphaMissense predictions of CFTR pathogenicity

AM makes pathogenicity predictions based on a 90% accuracy against ClinVar data^21^. For CFTR, AM predicted scores from 0.56 – 1.00 as pathogenic, scores from 0.34 – 0.56 as ambiguous, and scores from 0.04 – 0.34 as benign. We mapped the average AM prediction score per residue onto a CFTR structure (PDBID 5UAK)^25^ (**Figure 1A**). TMDs showed a propensity for pathogenicity in contrast to residue conservation as calculated by ConSurf^26^, which suggested the TMDs are comparatively variable across species (**Figure 1A & Supplemental Figure S1A**). Since the regulatory domain (RD) is disordered and not resolved in the CFTR structure 5UAK, we also plotted the average AM score for RD residues against a RD map highlighting key features such as transiently formed α-helices and phosphorylation sites^4^ (**Figure 1B**). Despite noted difficulty with disordered regions^21^, AM predicted RD residues ∼760-775 as a hotspot for pathogenicity. This is consistent with the role of transient helix 752-778 in CFTR gating through interactions with a conserved region of the NBD2 C-terminus^27,28^.

**Figure 1.**
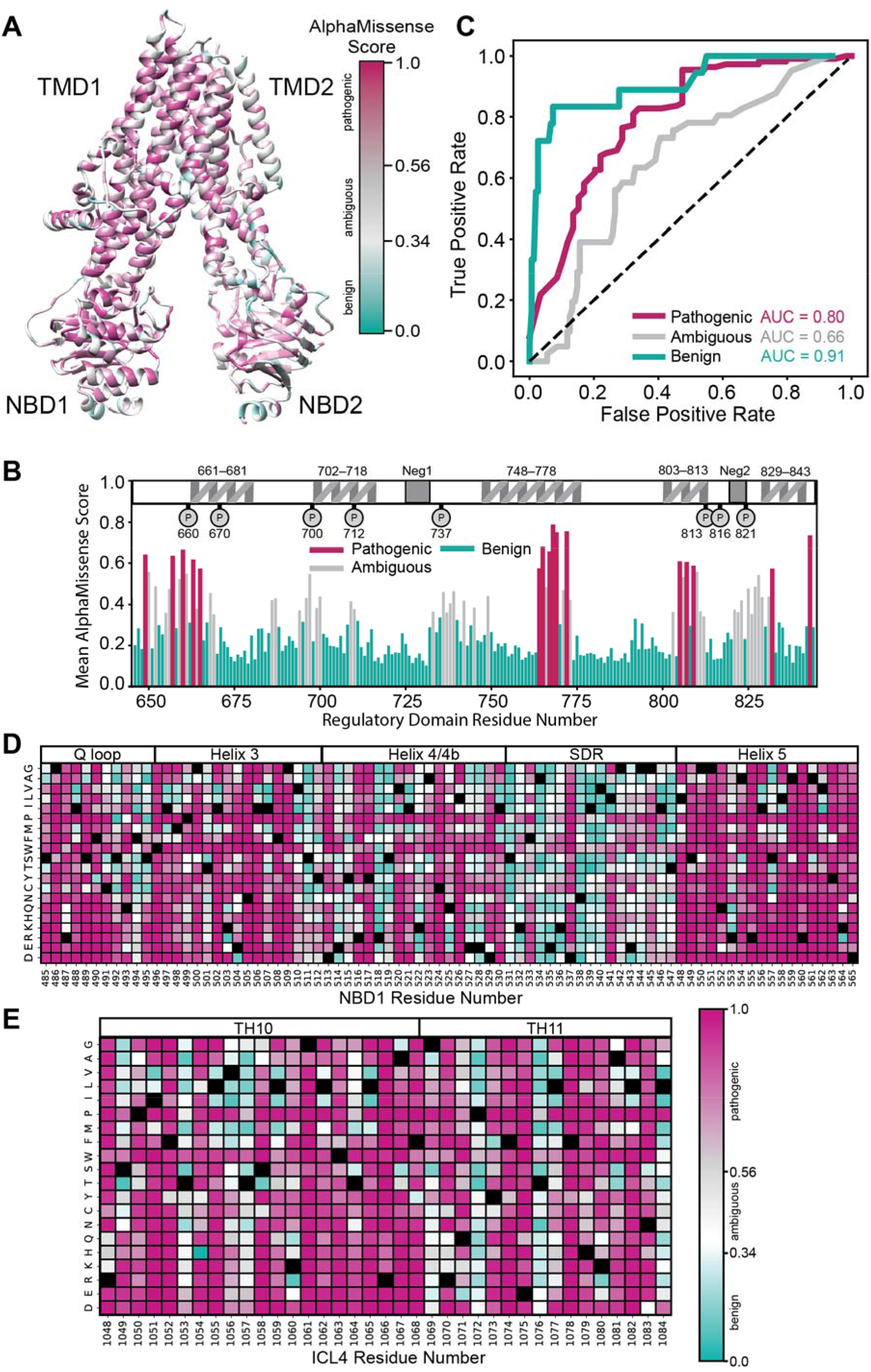
AlphaMissense predictions of CFTR pathogenicity compared to CFTR2.org repository. **A.** The average AM score per residue mapped onto the CFTR structure (PDBID 5uak)^4^. Variants with a score from 0.56-1.00 were classified by AM as pathogenic, variants with a score from 0.34 - 0.56 were classified as ambiguous, and variants with a score from 0.04 – 0.34 were classified as benign. **B.** The average AM score for the regulatory domain (RD) with an RD map of important features for reference^4^. Transient helices shown, two negatively charged regions (Neg1/2), and important phosphorylation sites shown as P. Notably, the second half of the transient helix 748-778 is predicted to be a hotspot of RD variant pathogenicity. **C.** Receiver operating characteristic (ROC) curve of AM predictions benchmarked against 169 CFTR2.org database classifications. Our curated data set contained 110 CF-causing, 41 variants variable clinical consequence (VVCC), and 18 non-CF causing missense variants. In the pathogenic curve (violet) – pathogenic prediction of a CF-causing variant was considered a true positive. Likewise in the ambiguous curve (grey) – ambiguous prediction of a variable clinical outcomes was considered a true positive. Finally, in the benign curve (bluegreen) – benign prediction of a non-CF-causing variant was considered a true positive. **D.** Heatmap of AM scores for NBD1 residues 485-565 including all regions interfacing with ICL4 and adjacent regions. Pathogenicity colored as in **A.** and WT residues depicted in black. **E.** Heatmap of AM scores for ICL4 residues 1048-1084, colored as in **E.**

We sought to evaluate AM’s ability to correctly predict CF pathogenesis on 169 classified missense variants from CFTR2.org^19^. The CFTR2 database offered a rich patient metric repository including pathogenicity classifications as CF-causing, VVCC, non-CF-causing, or VUS (**Supplemental Table 1**). VVCC are defined by CFTR2 as variants that may cause CF when found heterozygous with CF-causing variants, which results in variable clinical diagnosis of CF, e.g., a person with a VVCC and a CF-causing variant may or may not present with CF^19^.

AM showed a 95% accuracy (104/110) for predicting pathogenic variants and a 78% accuracy (14/18) for predicting benign variants based on variant determination in CFTR2 (**Supplemental Table 1**). We calculated the receiver operating characteristic (ROC) curve for all pairwise comparisons of pathogenicity predicted by AM (**Figure 1C, See Methods**). Briefly, all pairwise comparisons were considered – pathogenic, ambiguous, or benign were taken in turn to be a true positive. The alternative two predictions for a specific comparison were taken to be false positives. We considered pathogenic to predict CF-causing, ambiguous to predict VVCC, and benign to predict non-CF causing, VUS were not used. While looping through all possible score thresholds, the corresponding true positive and false positive rates were calculated and plotted. Benign predictions showed the highest area under the curve (AUC) (0.91) followed by pathogenic (0.80) and ambiguous respectively (0.66) – suggesting that AM has a high false positive rate, particularly for ambiguous predictions (**Figure 1C**). A high false positive rate may be attributed to a poor AlphaFold2 (AF2) predicted structure of CFTR. However, the AF2 predicted CFTR^29^ shows a root mean squared deviation of just 2.5 Å from the active state cryo-EM model (PDB ID 6MSM, resolution 3.2 Å^30^) (**Supplemental Figure S1B**).

We noted seven VUS in the CFTR2.org database and their respective AM predictions (**Table 1**). The location of these variants in the CFTR structure is shown (**Supplemental Figure S1C)**. Benign predicted R31L disrupts the arginine framed tripeptide motif at R29-R31, important for folding evaluation prior to ER export^31^ and may affect endocytosis rates^32^. V201M was ambiguously predicted, consistent with our previous report describing this variant as mildly mis-trafficked and selectively sensitive to VX-661^23^. A439V (benign prediction) and Y1014C (ambiguous prediction) showed trafficking and function slightly below WT^33^ suggesting these variants are benign. Benign predicted variant S912L lies close to the CFTR glycosylation sites at N894 and N900, thus we speculated this mutation could interfere with glycan processing. Nevertheless, S912L trafficking and function remained sufficient compared to WT *in vitro*^34^.

**Table 1.** CFTR2.org Variants of Unknown Significance (VUS). Seven missense variants of unknown significance (VUS) from the CFTR2.org database with their respective AM scores and predicted pathogenicity. Variants D923N and M952T are predicted to be pathogenic.

| Variant | AlphaMissense Score | Pathogenicity Prediction |
| --- | --- | --- |
| R31L | 0.17 | benign |
| V201M | 0.36 | ambiguous |
| A349V | 0.26 | benign |
| S912L | 0.12 | benign |
| D924N | 0.83 | pathogenic |
| M952T | 0.85 | pathogenic |
| Y1014C | 0.37 | ambiguous |

Variants D924N and M952T, both located in transmembrane helix 8, are predicted as pathogenic (**Supplemental Figure S1C)**. D924N resides in the potentiator binding hotspot^35,36^ and, according to clinical data, may cause pancreatic insufficiency but not lung disease^37^. M952T displays robust functional expression *in vitro*^38^, and two patients with an M952T/F508del genotype exhibit normal chloride transport measured from intestinal mucosa^38^ – suggesting this variant is likely not pathogenic, despite the AM prediction.

For performance comparison, we also plotted ROC curves for AM predictions of the 115 ClinVar variants from the AlphaMissense study and observed 96% average accuracy as presented previously^21^ (**Supplemental Figure S1D**). To validate this finding, we additionally downloaded a dataset of 209 ClinVar variants directly from ClinVar, including 96 overlapping variants from the AlphaMissense benchmark set. The ROC curve for our expanded ClinVar dataset showed >90% prediction accuracy with additional variants (**Supplemental Figure S1E**). Finally, we plotted a ROC curve for 113 ClinVar variants not included in the AM benchmark set, which revealed a >90% accuracy and indicates AM performs well on ClinVar data outside of the training set (**Supplemental Figure S1F-G**). In addition to classified variants used for performance evaluation, ClinVar contains 1,277 CFTR VUS^24^. AM predicted VUS ClinVar variants to contain 728 benign, 181 ambiguous, and 368 pathogenic variants (**Supplemental Figure S1H, Supplemental Table 2**).

AM performance was also compared to two other pathogenicity prediction tools, Evolutionary Scale Modeling (ESM)^39^ and Evolutionary model of Variant Effect (EVE)^40^ (**Supplemental Figure S2A-B**). In the ROC AUCs of benign variants, the AM value (0.91) was higher than those obtained for ESM (0.78) or EVE (0.78). A similar observation was made for pathogenic ROC AUCs, with AM (0.80) slightly above ESM (0.76) or EVE (0.73). ROC AUCs for ambiguous variants were nearly uniform across all methods (AM, 0.66; ESM, 0.65; EVE, 0.64). AM therefore offers a slight advantage for predicting pathogenic or benign variants and less utility regarding ambiguous variants.

Previous analysis of CFTR variants across sampled genetic information indicates the NBD1-intracellular loop 4 (ICL4) interface is a hotspot of pathogenicity^41^. Thus, we generated a heatmap of AM scores for NBD1 residues 485-565, which encompass the α-helical subdomain, structurally diverse region (SDR), and the entire NBD1-ICL4 boundary (**Figure 1D**). For the Q-loop (residues 486-495) and helix 3 (residue 496-512), potential substitutions are largely predicted as pathogenic except for residues 494 and 511. Of note, AM predicts position 508 as intolerant to substitution. Deletion of the encoded phenylalanine (F508del) is the most frequently reported variant among worldwide CF populations^8,42–44^. Most variations calculated as benign or ambiguous occur within helix 4/4b of the α-helical subdomain (residues 511-532) or SDR (residues 533-547) (**Figure 1D**). Possible substitutions across helix 4/4b that are predicted as pathogenic include V520 and C524. V520F^7^ and C524X^45^ are not presently approved for CFTR correctors and potentiators. Most substitutions (40% benign, 35% pathogenic) within the SDR are predicted as benign as expected base on the lack of structure in the region.

In contrast, variations at the NBD1-ICL4 interface are overwhelmingly scored as severe. Residues 548-565 comprise the NBD1 core helix 5, which directly interacts with ICL4 and demonstrates the strongest sensitivity (7% benign, 82% pathogenic) to mutation with potential substitutions predicted as pathogenic (**Figure 1D**). This region contains numerous CF-causing variants, some of which are refractory to available CFTR modulators, such as R560T/K/S^4,41^. Within the ICL4 region (residues 1048-1084), AM scores indicate 14% benign and 69% pathogenic predictions (**Figure 1E**). The heatmap reveals residues 1069, 1072, 1076, and 1084 as relatively tolerant to substitution. Together, these data suggested AM pathogenicity scores matched previous findings, as well as our general understanding about residue conservation throughout CFTR, while providing specific information about every possible substitution.

### II. Cystic Fibrosis pathogenicity correlated modestly with AlphaMissense predictions

In addition to classifying variant pathogenicity, the CFTR2.org database annotates clinical outcomes for persons with CF including sweat chloride levels, pancreatic insufficiency rates, *Pseudomonas aeruginosa* infection rates, and lung function^19^. We curated the clinical outcomes for all CFTR missense variants with available data (**Supplemental Table 1, See Methods**). We then analyzed the ability of AM to predict patient pathogenicity metrics. Briefly, CFTR2.org data were downloaded from the Variant List History tab and filtered for 176 missense variants (169 classified and 7 VUS). Then, clinical outcome data were manually assembled by searching each variant and recording the sweat chloride (mEq/L), pancreatic insufficiency rate (%), *P. aeruginosa* infection rate (%), and lung function (forced expiratory volume in one second (FEV1), % predicted). Of note, CFTR2 data was based on individual alleles, e.g. missense variants.

First, we plotted AM score versus CF sweat chloride levels for 123 missense variants with sweat chloride values reported (**Figure 2A**). AM score correlated modestly with sweat chloride levels (Pearson Correlation Coefficient: 0.46, Spearman Correlation Coefficient: 0.48). CF-causing variants, shown in blue, clustered in the top right corner, indicative of high AM scores and elevated sweat chloride levels. By contrast, VVCCs, shown in yellow, clustered in the bottom right corner, reflecting an excessive AM score (**Figure 2A**). When considering CF-causing or VVCC separately, we note a reduced correlation between sweat chloride levels and AM scores (**Supplemental Figure 3A-B**), suggesting AM captures the trend across all variant types rather than performing better on pathogenic variants.

**Figure 2.**
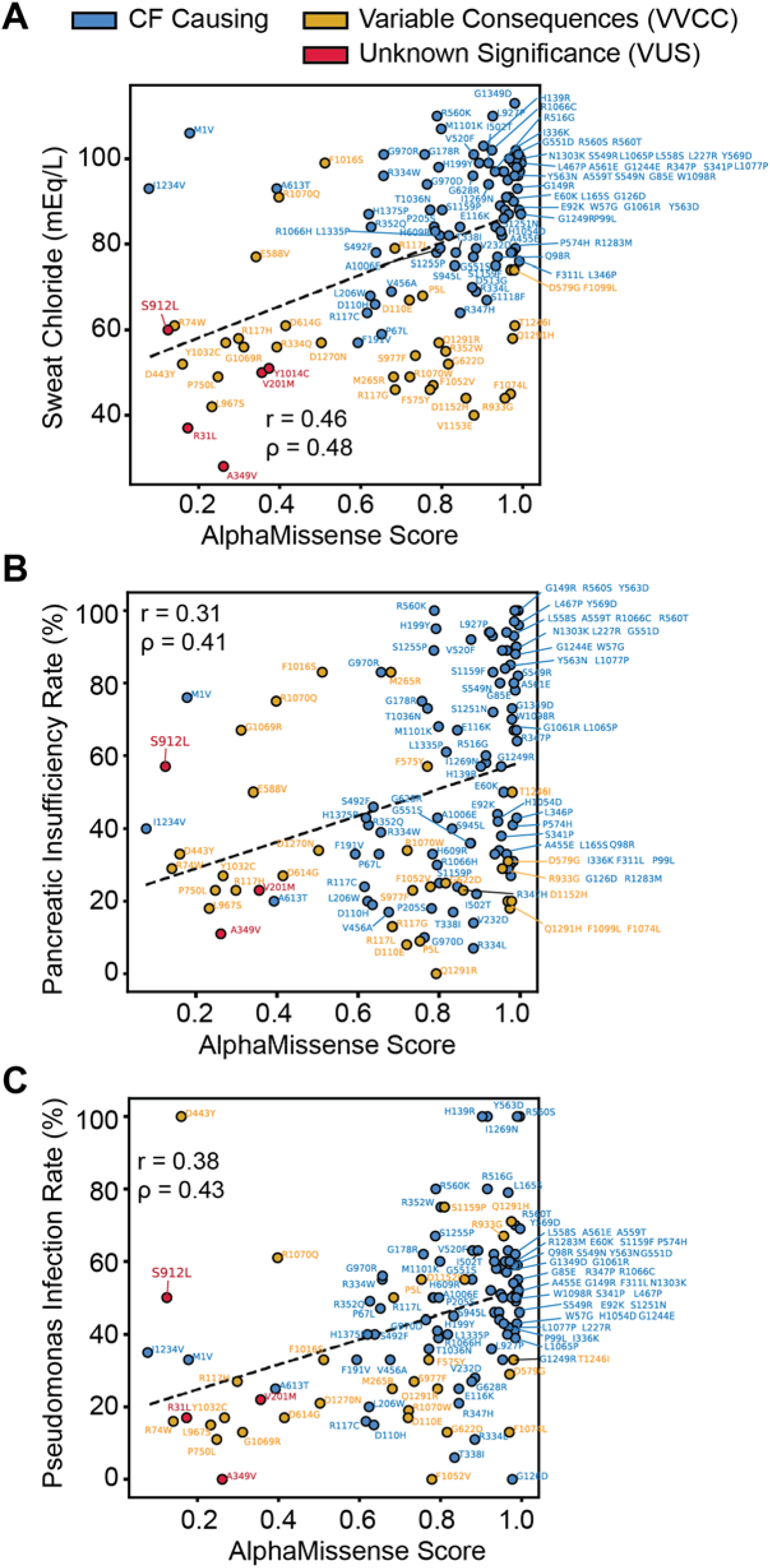
Benchmarking AlphaMissense against Cystic Fibrosis patient pathogenicity metrics. **A.** AM score plotted against sweat chloride levels in milliequivalents per liter (mEq/L) for 123 missense variants. Healthy sweat chloride levels were <30 mEq/L. CF-causing variants were shown in blue, variants of variable clinical consequence (VVCC) were shown in yellow, and variants of unknown significance (VUS) were shown in red (as annotated in CFTR2). A modest linear correlation (Pearson Coefficient r = 0.46, Spearman Coefficient ρ = 0.48) was observed. **B.** AM score plotted against pancreatic insufficiency rates in percent for 116 missense variants. Less correlation (Pearson Coefficient r = 0.31, Spearman Coefficient ρ = 0.41) was observed than with sweat chloride, notably many VVCCs were predicted pathogenic AM score but demonstrate a low pancreatic insufficiency rate. Colors annotated as in **A.** **C.** AM score plotted against *P. aeruginosa* infection rates in percent for 114 missense variants. Colors annotated as in **A.** Linear correlation (Pearson Coefficient r = 0.38, Spearman Coefficient ρ = 0.43) is shown. Interestingly, S912L, a variant of unknown significance is predicted to be benign but shows a high sweat chloride, pancreatic sufficiency, and *P. aeruginosa* infection rate compared to variants with similar scores.

Next, we plotted AM score versus pancreatic insufficiency rates for 116 missense variants present on at least one allele of persons with CF with CFTR2 outcomes reported (**Figure 2B**). AM scores correlated poorly with pancreatic insufficiency rates (Pearson coefficient: 0.31, Spearman Coefficient: 0.41) compared to sweat chloride. Again, AM failed to predict VVCCs, shown in yellow, on this metric (**Figure 2B**). However, considering CF-causing and VVCCs separately failed to change the correlation for pancreatic insufficiency (**Supplemental Figure 3C-D**). Finally, we plotted AM score versus *P. aeruginosa* infection rates for 114 missense variants on at least one allele with CFTR2 outcomes reported (**Figure 2C**). AM correlated better here than for pancreatic insufficiency rates, but worse than for sweat chloride (Pearson Coefficient: 0.38, Spearman Coefficient: 0.44). However, it performed better on VVCCs, yet correlation was again reduced when only CF-causing or VVCCs were separately considered (**Supplemental Figure 3E-F**).

Taken together, AM correlated modestly with clinical data and performed poorly on VVCCs and VUSs. For example, VUS S912L was predicted benign with an AM score of 0.12. However, this variant was associated with sweat chloride levels of 60 mEq/L (**Figure 2A**), which resides exactly at the diagnostic cutoff for CF. S912L displays a pancreatic insufficiency rate of 57% (**Figure 2B**) and *P. aeruginosa* infection rate of 50% (**Figure 2C**) – suggesting this variant may present with more pathologic characteristics than predicted or annotated in CFTR2. Unfortunately, pathogenic-predicted variants such as D924N and M952T have insufficient data available on CFTR2 for comparison. Weak performance by AM could be attributable to high false positive rates and/or compound heterozygous genotypes. The latter factor likely complicates interpretation of clinical data, as people with complex CF alleles may exhibit differing degrees of variant severity on each chromosome (e.g. one CF-causing paired with a VUS/VVCC) compared to patients with the same variant severity on each allele (e.g. two CF-causing).

### III. AlphaMissense predicts CFTR function beyond folding and trafficking competency

Much CFTR biochemical and functional data was also available for comparison, including recent deep mutational scanning (DMS), theratype screening, and spatial covariance studies^23,33,46^. In the DMS study, fluorescence-activated cell sorting was used to measure the cell surface immunostaining intensity of an epitope-tagged library of 129 CFTR variants including 100 missense variants^23^. In the theratype study, 655 variants including 585 missense variants were screened for their trafficking efficiency and function^33^. In the spatial covariance study, a CFTR trafficking and a chloride conductance index were established to characterize variant temperature response^46^. Variable, albeit high, overlap was observed between the CFTR2 dataset and the *in vitro* data sets discussed below (**Supplemental Figure S2C**).

First, we evaluated AM ability to predict CFTR folding competency – which is well characterized to correlate with cell surface expression and trafficking efficiency^47–50^. We plotted AM prediction scores for 100 missense variants versus DMS cell immunostaining intensity (**Figure 3A**), which showed an inverse relationship with poor correlation (Pearson coefficient: -0.37, Spearman Coefficient: -0.37). Notably, among variants in the top right corner, e.g. high AM score and high surface staining, we observed several gating variants (G551D/S, R347H, S1251N, and G1244E etc.) (**Figure 3A**). Mis-gating variants traffic normally, but they are CF-causing due to disrupted properties of channel opening and closing. This result demonstrated that AM failed to infer the nature of the variant defect.

**Figure 3.**
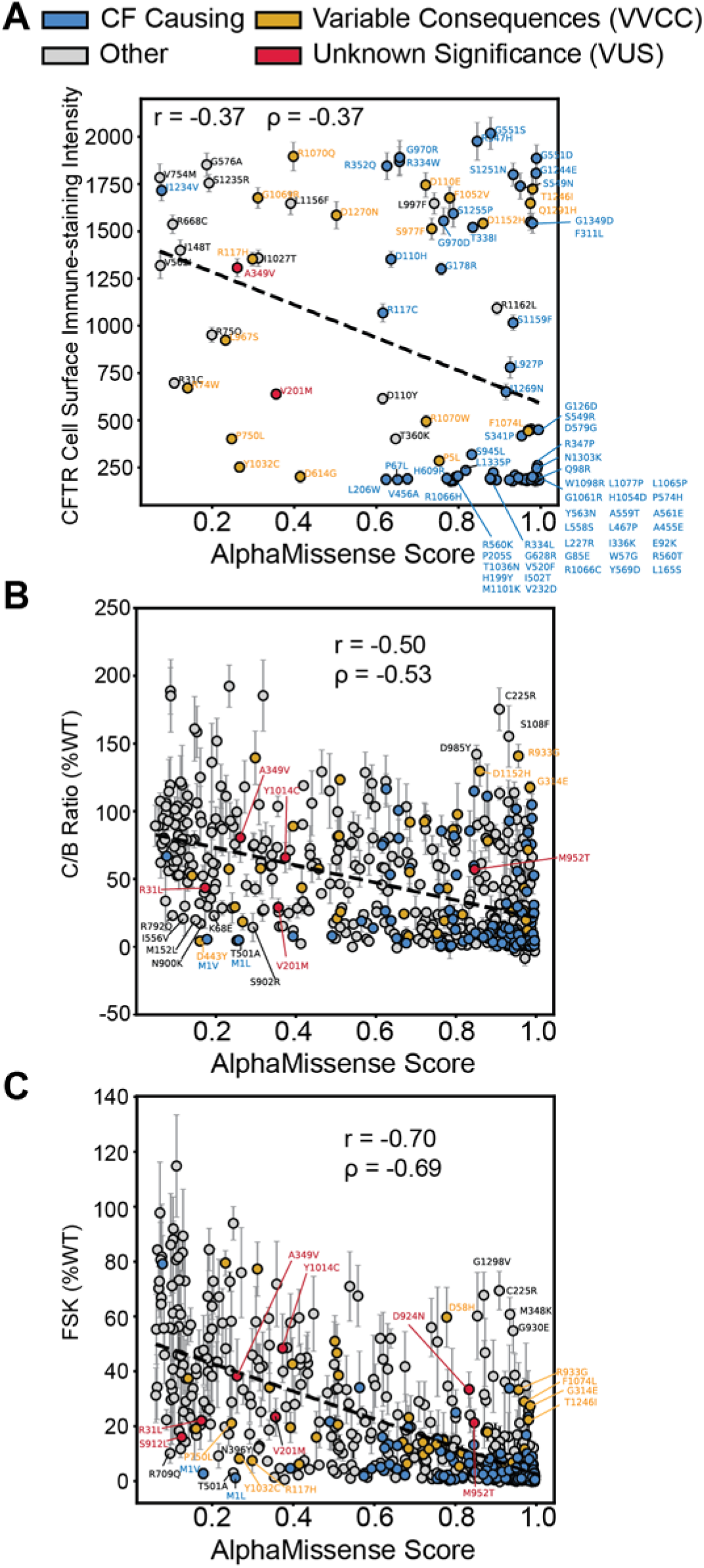
Benchmarking AlphaMissense against CFTR *in vitro* functional metrics. **A.** AM score plotted against deep mutational scanning data for 100 missense variants in HEK293T cells^23^. The y-axis represents the cell surface immune-staining intensity of CFTR and thus is indicative of the CFTR trafficking levels to the cell surface. A slight inverse linear correlation was observed (Pearson Coefficient r = - 0.37, Spearman Coefficient ρ = -0.37). Some off-axis variants such as G551D are gating variants and thus fail to experience aberrant trafficking in the HEK293T cell background but still exhibit impaired channel function. CF-causing variants are shown in blue, variants of variable clinical consequence (VVCC) are shown in yellow, and variants of unknown significance (VUS) are shown in red (as annotated in CFTR2). **B.** AM score plotted against CFTR western blot C:B band ratio in percent WT for 538 missense variants^33^. Several off-axis variants are highlighted. Color annotation as in **A.** Inverse linear correlation (Pearson Coefficient r = -0.50, Spearman Coefficient ρ = -0.53) improves for this larger dataset. Of note, VUS D924N and S912L were filtered out of this dataset due to an SEM >30, indicative of poor reproducibility across replicates (**Supplemental Figure 4A-B**). **C.** AM score plotted against Forskolin induced CFTR current in percent WT for 546 missense variants^33^. Color annotation as in **A.** Inverse linear correlation (Pearson Coefficient r = -0.70, Spearman Coefficient ρ = -0.69) is higher than the function data compared to trafficking alone. All seven CFTR2 VUSs are highlighted.

Next, we plotted AM prediction scores for 538 missense variants versus CFTR trafficking efficacy as measured by the ratio of mature, fully-glycosylated CFTR (band C) to the immature glycoform (band B) on western blot (C/B band ratio) (**Figure 3B**)^33^. Experimental data was filtered for plotting clarity (**Supplemental Figure 4, See Methods**). We removed highly variable experimental data with a standard error of the mean (SEM) greater than 30. Most CFTR variants show a C:B ratio less than 30% of WT, indicating a lack of reproducibility for these measurements with higher variability (8% of data points removed, 92% retained). AM scores displayed improved inverse correlation with the larger trafficking efficiency dataset (Pearson coefficient: -0.50, Spearman Coefficient: -0.53). This finding suggested AM can predict CFTR folding competency across diverse types of variants. Several off-axis variants were annotated that show poor predictions and poor trafficking (<30% of WT) based on the distribution of all trafficking data (**Supplemental Figure 4A-B**).

Finally, we evaluated AM ability to predict CFTR function as measured by transepithelial current clamp conductance^33^. We plotted AM prediction scores versus forskolin (FSK)-induced basal CFTR channel activity as percent WT (FSK %WT) for 546 missense variants (**Figure 3C**). Again, highly variable experimental data were filtered out considering an SEM greater than 20 as most variants were less than 20% of WT (**Supplemental Figure 4, See Methods**), leaving 93% of the experimental data for comparison to AM. AM scores inversely correlated best with CFTR function measured by conductance (Pearson coefficient: -0.70, Spearman Coefficient: -0.69). Several off-axis variants were noted which show poor predictions and poor channel function (<30% of WT) based on the distribution of functional data (**Supplemental Figure 4C-D**).

We verified the increased capability to predict CFTR function by correlating AM scores with a spatial covariance study (**Supplemental Figure S5**). This study describes trafficking (measured by western blot band shift assay) and chloride conductance indices and presents data for both metrics at 37 °C and reduced temperature (27 °C)^46^. Reduced temperature is a well-established method for partially rescuing F508del biogenesis^51^. We observed a modest correlation (Pearson coefficient: -0.46, Spearman Coefficient: -0.44) with trafficking index at 37 °C, and a similar correlation at 27 °C (Pearson coefficient: -0.48, Spearman Coefficient: - 0.49) (**Supplemental Figure S5A-B**). Again, correlation increased when compared to chloride conductance index (Pearson coefficient: -0.58, Spearman Coefficient: -0.54 at 37 °C vs. Pearson coefficient: -0.50, Spearman Coefficient: -0.53 at 27 °C) (**Supplemental Figure S5C-D**). Together these results indicated that AM scores are closely aligned with pathogenicity but cannot differentiate between variants that compromise expression versus function.

### IV. AlphaMissense cannot predict CFTR variant theratype

Given the rapid and continuous emergence of novel CFTR variants detected by next-generation sequencing technologies, as well as a robust pipeline of new modulators and other CFTR-directed treatments under development, the need remains for optimized approaches to CF precision therapeutics. CFTR variant theratyping is an established method for quantifying *in vitro* CFTR sensitivity to pharmacologic agents, results of which are utilized to predict treatment responses for genotype-matched patients^6,52^. CF treatment involves two corrector compounds, VX-661 and VX-445, that likely bind directly to two unique sites on CFTR^14^, show distinct mechanisms, and hence distinct response profiles across variants. Thus, theratyping variant response remains an important task for CF personalized medicine.

We sought to determine whether AM offered any predictive power for CFTR theratyping, although this task is beyond the intended scope of AM. Theratype distinguishing plots were generated and colored by AM pathogenicity score to assess for potential patterns. We split VX-445-sensitive variants from VX-661-sensitive variants along a diagonal axis of best fit by plotting CFTR immunostaining intensity for VX-445 versus VX-661 (**Figure 4A**). Variants responsive to VX-445 fell above the dotted line, and variants responsive to VX-661 fell below the dotted line^23^. Variants were then colored by AM pathogenicity score, although the color distribution across the responsive spectrum revealed little discernable patterns (**Figure 4A**). We also plotted basal CFTR immune staining intensity versus VX-661, VX-445, or the combination thereof, then shaded variants by AM pathogenicity (**Supplemental Figure S6A-C**). Similarly, AM scores showed little-to-no color patterns and appear randomly distributed.

**Figure 4.**
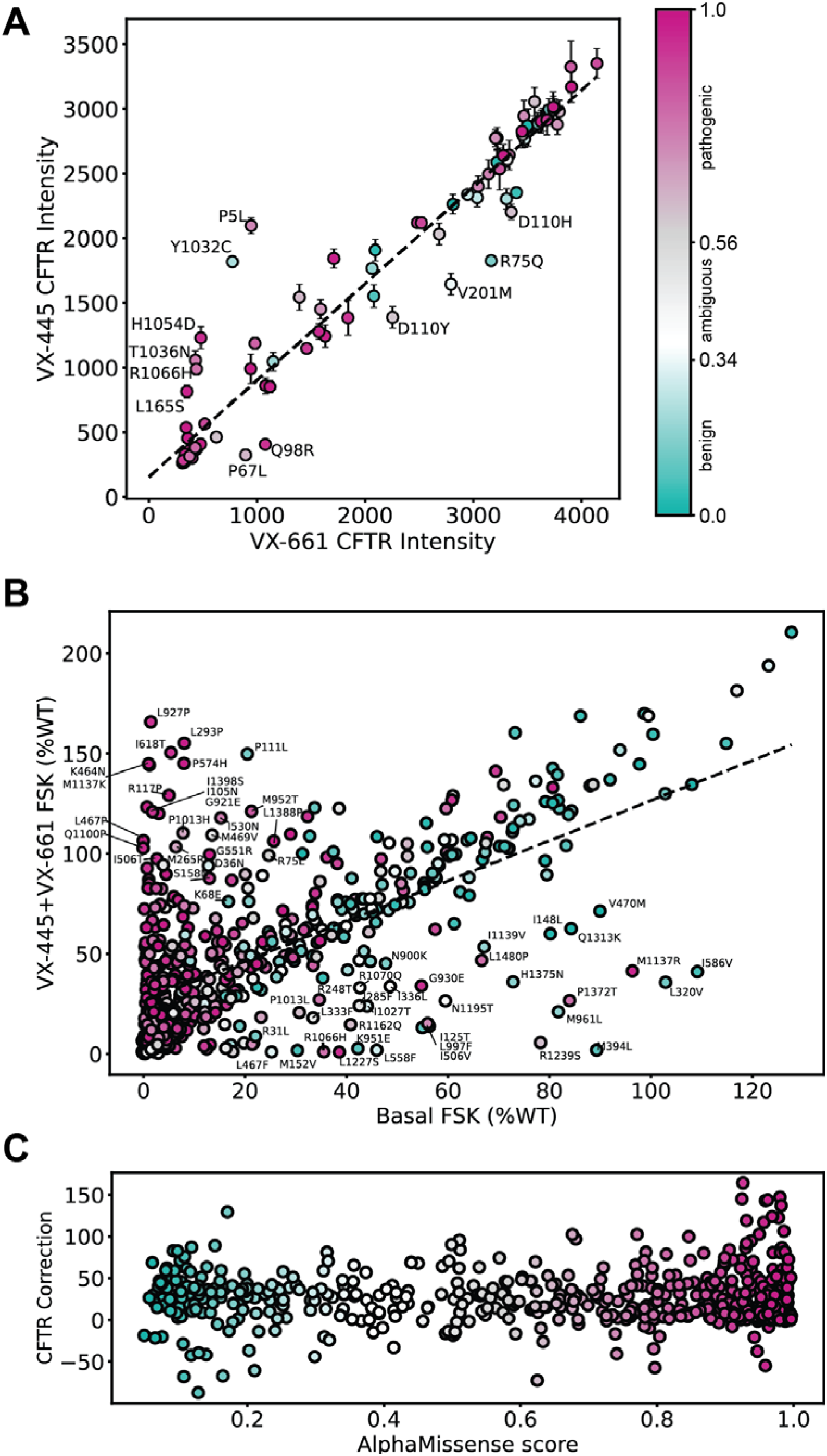
CFTR theratype plots colored by AlphaMissense pathogenicity score. **A.** CFTR cell surface immune staining intensity comparing treatment with VX-661 versus VX-445 correctors with the dotted line representing equivalent response to both correctors^23^. Variants that fell below the best-fit dotted line are selectively responsive to VX-661 while variants above the dotted line were selectively responsive to VX-445. The AlphaMissense pathogenicity predictions (color gradient) show no correlation with the corrector response patterns for CFTR variants. Variants with a score from 0.56-1.00 were classified by AM as pathogenic (violet), variants with a score from 0.34 - 0.56 were classified as ambiguous (grey), and variants with a score from 0.04 – 0.34 were classified as benign (green). Error bars represent the standard deviation of the cell surface immunostaining intensity. Selectively sensitive variants were annotated.

Next, we used the theratyping study CFTR functional data^33^ to plot the VX-445 + VX-661 FSK-mediated response (% of WT) versus basal activity, then colored the values by AM pathogenicity score (**Figure 4B**). Benign variants fell along a linear diagonal, suggesting that benign predicted variants all experience a linear response to CFTR correctors. We speculate this shift may reflect well-documented WT modulator response, implying an inherent stabilizing effect of VX-445 and VX-661. C:B band ratio response colored by AM score portrayed a random distribution of score color (**Supplemental Figure 7A**). Pathogenic predicted variants in both plots show a random distribution. To determine whether theratype was predicted by variant structural location within CFTR, combined with AM score, we subdivided the plot in **Figure 4B** by domain (**Supplemental Figure 7B-E**). Each domain individually showed a similar random distribution of score colors. Finally, we calculated relative degree CFTR correction by subtracting basal FSK (% of WT) from VX-445+VX-661 correction FSK (% of WT) and plotted this difference against AM score (**Figure 4C**). Again, no obvious pattern was observed. In summary, we found AM score afforded little predictive power for profiling pharmacologic responsiveness of CFTR variants. However, AM score could potentially be a useful machine learning feature for future theratype prediction methods.

## Conclusion

AlphaMissense has the exciting potential to aid with pathogenicity classification of rare and emerging variants identified during genetic screening. CF posited a valuable case study for evaluating AM performance because of abundance of clinical outcome data and *in vitro* variant classifications available. AM predicted pathogenicity of severe CF-causing variants well, albeit with a high false positive rate, and matched previous studies of CFTR variant pathogenicity in the NBD1/ICL4 interface^41^. However, AM performed modestly for pathogenicity predictions of VUSs and VVCCs, and the tool does not appear useful for CFTR theratype predictions. Again, for pathogenic missense variants, AM score correlated modestly with trafficking data and correlated well with channel activity functional data. Thus, predictions offer little information for distinguishing pathogenicity mechanism. AM may provide guidance in determining if polymorphisms in CFTR are benign, but performance on less severe disease variants indicate that caution must be taken when interpreting AM predictions. *In vitro* measurements on variant severity may aid in evaluating prediction quality and will remain necessary for CFTR theratyping.

## Methods

### Data Curation and Collection

AlphaMissense (AM) predictions for all single amino acid substitutions in the human proteome data was downloaded, gunzipped, and searched using vim text editor for CFTR accession number/Uniprot ID P13569. CFTR AM predictions were extracted into a separate file for analysis. ESM score predictions were downloaded from https://huggingface.co/spaces/ntranoslab/esm_variants by searching accession number P13569. EVE predictions were downloaded from https://evemodel.org/proteins/CFTR_HUMAN#variantsTableContainer by searching accession number P13569.

Cystic Fibrosis clinical outcome data was initially downloaded from the Variant List History tab on CFTR2.org. The table of 804 variants was filtered for 176 missense variants by removing in/dels, splicing variants, premature stop codons, etc. The patient information was manually curated by searching each variant and annotating the sweat chloride (mEq/L), pancreatic insufficiency rate (%), *P. aeruginosa* infection rate (%), lung function ages < 10 (FEV1%), lung function ages 10<20 (FEV1%), lung function ages >20 (FEV1%) (**Supplemental Table 1**). Lung function data proved too highly variable for comparison and was not used, but was still included in the Supplemental Table for reference. CFTR2 definitions for these variants are as follows^19^: CF-causing: “A variant in one copy of the *CFTR* gene that always causes CF, as long as it is paired with another CF-causing variant in the other copy of the *CFTR* gene.” Non CF-causing: “A variant in one copy of the *CFTR* gene that does not cause CF, even when it is paired with a CF-causing variant in the other copy of the *CFTR* gene.” Variant of Variable Clinical Consequence (VVCC): “A variant that may cause CF, when paired with a CF-causing variant in the other copy of the *CFTR* gene.” Variant of Unknown Significance (VUS): “A variant for which we do not have enough information to determine whether or not it falls into the other three categories.”

*In vitro* modulator response data was downloaded from^33^ and deep mutational scanning data downloaded from^23^. The 650 variants from^33^ were filtered to 585 missense variants. ClinVar data was downloaded after searching for CFTR. Clinvar predictions were filtered for missense variants by removing, in/dels, stop codons, double missense variants, etc. Filtering yielded 1768 missense variant pathogenicity predictions (**Supplemental Table 2**). For performance comparison, the missense variants were filtered by clinical significance. We removed classifications such as *no interpretation*, *conflicting interpretations*, *uncertain significance*, and *drug response*. This left 219 variants classified as *pathogenic* or *likely benign* for performance evaluation and ROC plotting.

### Filtering Experimental Data

Experimental data from Bihler et al.^33^ were filtered to exclude highly variable data based on the SEM due to lack of reproducibility. We plotted both the distribution of the data itself to look for outliers on the y axis of our correlation plots (**Figure 3B-C**) and the distribution of the SEM (**Supplemental Figure 4**). We labeled outliers with a C:B ratio of less than 30, but with a benign AM prediction of less than 0.3 (**Supplemental Figure 4A, Figure 3B**). C:B band ratio SEM of greater than 30 were excluded from the analysis and not plotted for clarity, leaving 538 variants for analysis – 92% of the experimental data (**Supplemental Figure 4B**). For the functional data, we labeled outliers of interest with a FSK % of WT of less than 30, but with a benign AM prediction of less than 0.3 (**Supplemental Figure 4C, Figure 3C**). FSK % of WT SEM of greater than 20 were excluded from the analysis and not plotted for clarity, leaving 546 variants for analysis – 93% of the available experimental data.

### Analysis

Data were analyzed and plotted in Python 3. Raw excel files were imported and parsed using the Pandas data frame library and plots were generated with the matplotlib.pyplot and seaborn libraries. Pearson and Spearman correlation coefficients were calculated with the scipy.stats library using the pearsonr() and spearmanr() functions respectively. Plots were generated for all possible variants with available data for a given metric. The receiver operating characteristic (ROC) curve for AM pathogenicity predicted by AM was calculated against CFTR2 classification. CFTR2 classifies variants as CF causing, variable clinical consequence (VVCC), or non-CF causing. We equated the AM prediction pathogenic to CF causing, ambiguous to variable, and benign to non-CF causing. Since ROC is used for binary classification, all pairwise comparisons were considered. In each ROC curve, a different prediction (pathogenic, ambiguous, or benign) was taken to be a true positive, and the other two predictions to be false positives. Then the corresponding true positive and false positive rates were calculated by considering all possible score cutoffs for pathogenicity. Theratype discerning plots were generated to distinguish responsive from non-responsive variants graphically and colored by the variants respective AM score.

## Author Contributions

Conceptualization: E.F.M., L.P. Funding acquisition: E.F.M., K.E.O., J.P.S., L.P., J.M. Data Collection: E.F.M., K.E.O. Data Analysis: E.F.M. WritingLoriginal draft: E.F.M.; WritingLreview and editing, E.F.M., K.E.O., J.M., L.P. All authors have read and agreed to the published version of the manuscript.

## List of Supplemental Tables

Supplemental Table S1: CFTR2.org variant clinical outcome data

Supplemental Table S2: ClinVar CFTR variant data

Supplemental Table S3: In vitro CFTR trafficking and functional data from Bihler et al

Supplemental Table S4: Deep Mutational Scanning CFTR data

