## Supplemental Information for "Benchmarking AlphaMissense Pathogenicity Predictions Against Cystic Fibrosis Variants"

Table of Contents

Supplemental Figures........................................ 2-13

References........................................ 14

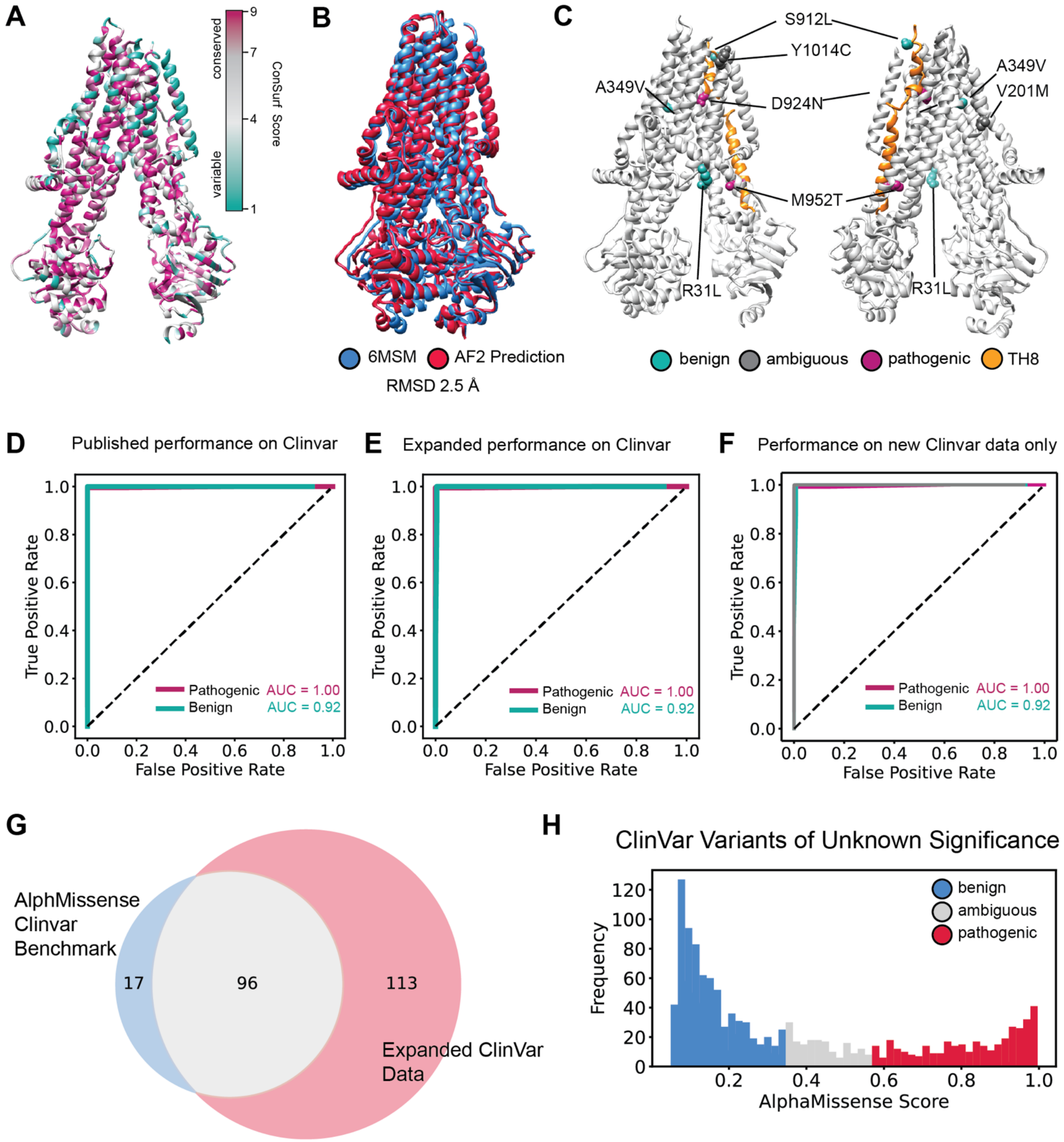


**Supplemental Figure S1. AlphaMissense prediction of CFTR variants of unknown significance (VUS)**

**A.** Conservation of residue in CFTR mapped on the structure (PDBID 5UAK)[1,2]. The abundance of green, representing low conservations scores in the TMDs stands in notable contrast the AM score predictions of pathogenicity in the TMDs.

**B.** An overlay of the active state CFTR (PDB ID 6MSM)[3] and the AlphaFold prediction for CFTR[4] showing nearly perfect alignment of all resolved residues (1-409,435-637, 845-889, 900-1173, 1202-1451). We calculated the root mean squared deviations (RMSD) of carbon backbone atoms between these two models in Chimera and found an RMSD of just 2.5 Å.

**C.** Variants of unknown significance (VUS) displayed on CFTR structure (PDBID 5UAK)[1] demonstrates all unknown variants are in the transmembrane domains. Benign predicted mutations are shown in green, ambiguous predicted mutations in grey, and pathogenic predicted mutations are shown in purple. The two pathogenic predicted mutations both occur in transmembrane helix 8 (TH8) shown in orange.

**D.** Receiver operating characteristic curve for AlphaMissense predictions of 115 Clinvar variants presented in the AlphaMissense benchmark. The average performance between pathogenic and benign variants is 95.8% as previously presented[5].

**E.** Receiver operating characteristic curve for AlphaMissense predictions of 209 variants downloaded directly from Clinvar[6], including 96 overlapping variants from the AlphaMissense benchmark. Again, the average performance is 95.8%.

**F.** Receiver operating characteristic curve for AlphaMissense predictions of 113 variants downloaded directly from Clinvar[6] that did not overlap with variants from the AlphaMissense benchmark. Despite, not being trained on these ClinVar data, average performance is 95.8%.

**G.** Overlap between AlphaMissense ClinVar benchmark set and our extended ClinVar set – showing 115 variants from AM and an additional 113 variants considered in **F.** Performance of AlphaMissense is very good across all permutations of ClinVar data considered.

**H.** Due to the high number of VUS predictions in ClinVar[6] for CFTR missense mutations, we plotted the AlphaMissense score for all 1277 VUSs in ClinVar. We show 728 benign, 181 ambiguous, and 368 pathogenic variants as predicted by AM. Data is available in Supplemental Table 2.

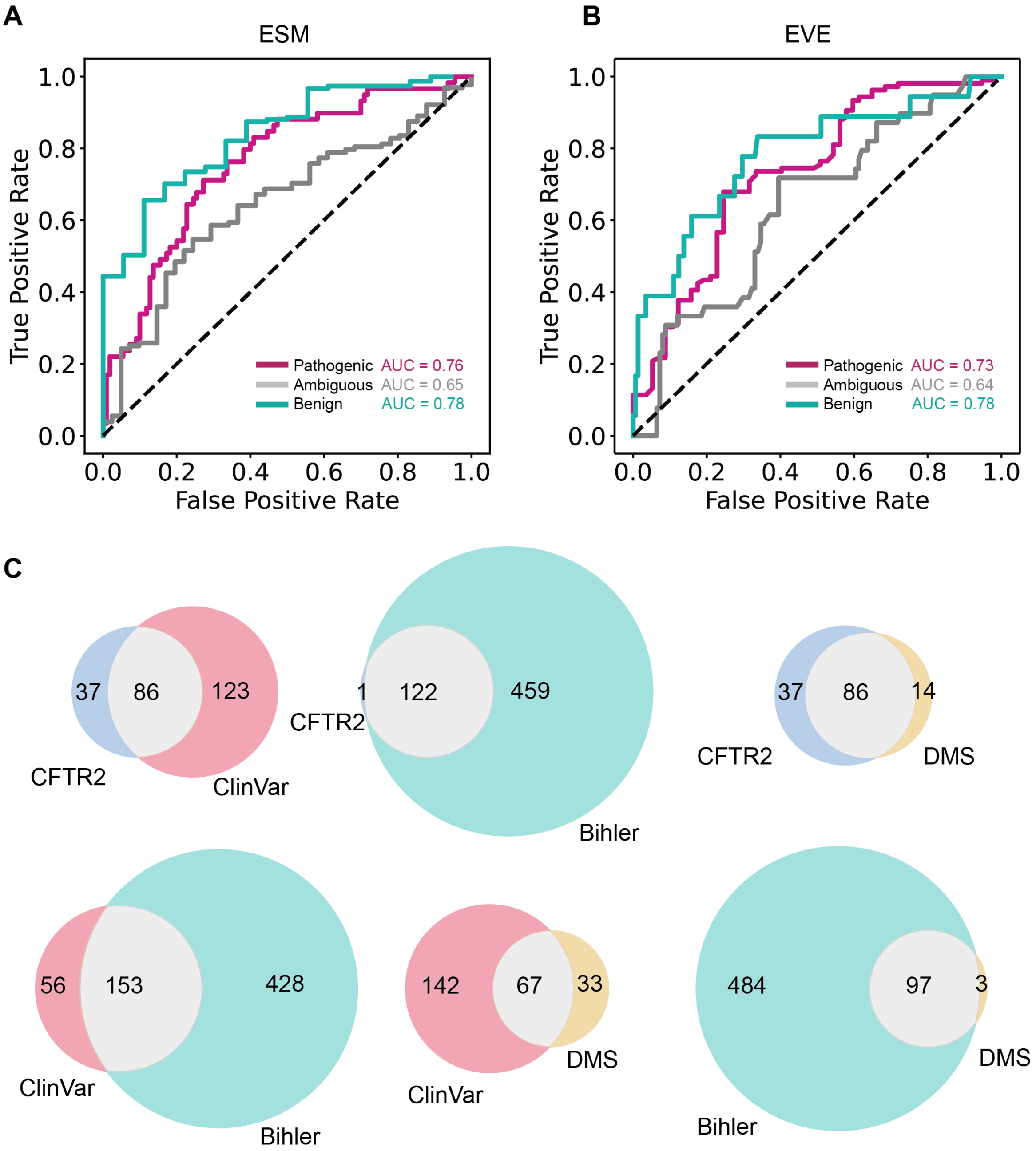


**Supplemental Figure S2. Alternative Prediction method performance and dataset overlap.**

**A.** Receiver operating characteristic curve for ESM predictions[7] of 169 CFTR missense variants including 110 CF causing, 41 variable clinical consequence (VVCC), and 18 non-CF causing variants. For the pathogenic curve (violet), we considered a pathogenic prediction of a CF-causing variant a true positive. For the ambiguous curve (grey) - we considered the ambiguous prediction a VVCC a true positive. For the benign curve (bluegreen) – we considered the benign prediction of a non-CF causing variant as a true positive.

**B.** Receiver operating characteristic curve calculated the same as in **A.** but using EVE missense variant predictions[8] of 169 CFTR missense variants colored as shown in **A**.

**C.** Venn diagrams depicting the overlap of various datasets used throughout the study. We considered our expanded ClinVar dataset, the deep mutational scanning (DMS) dataset[9], our curated CFTR2 dataset, and the missense variants from the Bihler et al. dataset[10].

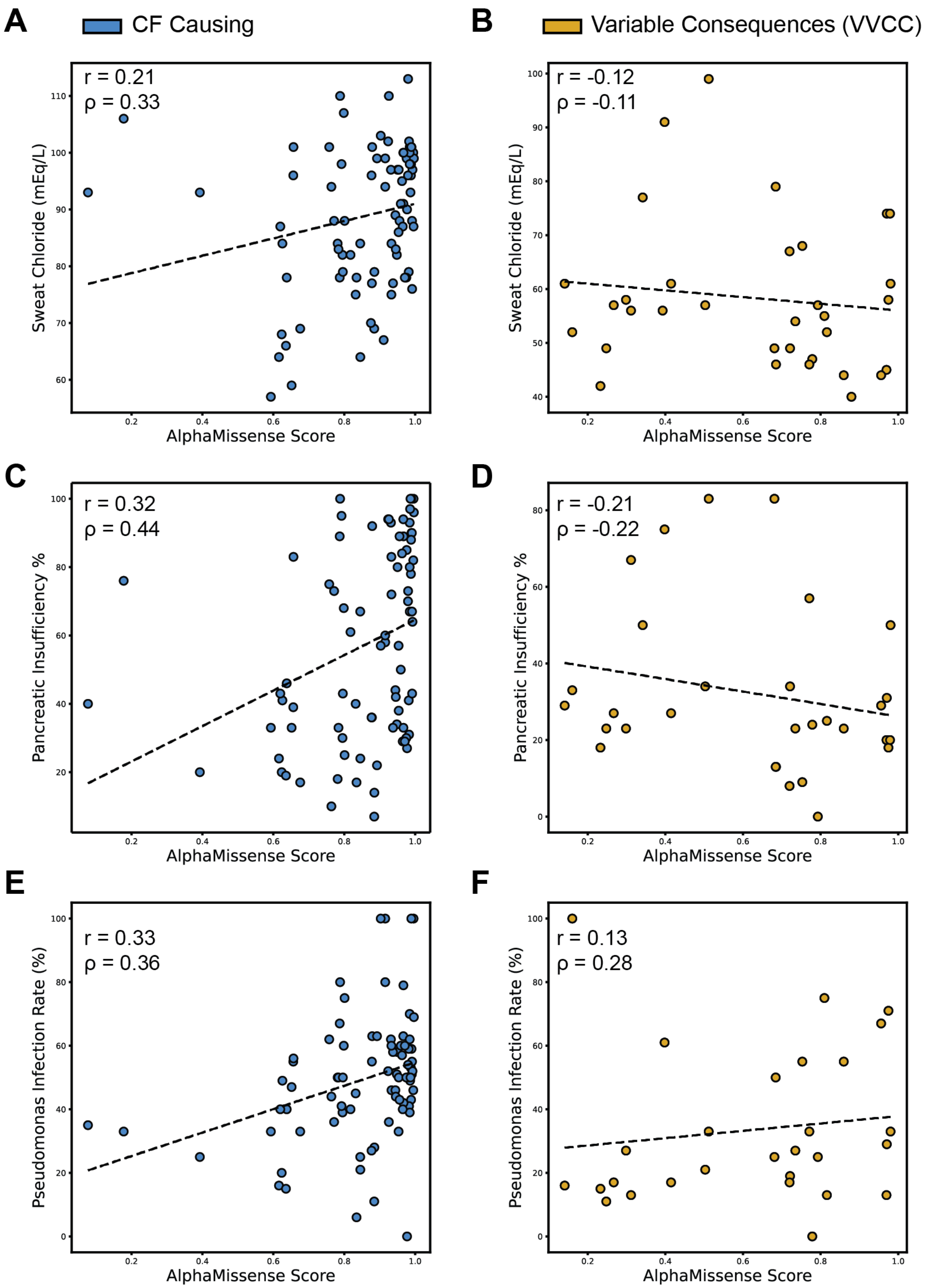


**Supplemental Figure S3. AlphaMissense prediction correlation s with Cystic Fibrosis patient pathogenicity metrics by diagnosis**

**A.** AM score plotted against sweat chloride levels in milliequivalents per liter (mEq/L) for 85 missense variants classified as CF causing. The linear correlation (Pearson Coefficient r = 0.21, Spearman Coefficient ρ = 0.33) is reduced compared to the complete data set correlation shown in **Figure 2A**.

**B.** AM score plotted against sweat chloride levels for 33 missense variants classified as variants of variable clinical consequence (VVCC). The linear correlation (Pearson Coefficient r = -0.12, Spearman Coefficient ρ = -0.11) is statistically insignificant.

**C.** AM score plotted against pancreatic insufficiency rates in percent for 83 missense variants classified as CF causing. The correlation (Pearson Coefficient $r = 0.32$, Spearman Coefficient $\rho = 0.44$) was similar to the entire dataset in **Figure 2B**.
**D.** AM score plotted against pancreatic insufficiency rates for 30 missense variants classified as VVCC. The correlation for these data (Pearson Coefficient $r = -0.21$, Spearman Coefficient $\rho = -0.22$) was statistically insignificant.
**E.** AM score plotted against pseudomonas infection rates for 82 missense variants classified as CF-causing. Linear correlation is reduced compared to the entire data set presented in **Figure 2C** (Pearson Coefficient $r = 0.33$, Spearman Coefficient $\rho = 0.36$).
**F.** AM score plotted against pseudomonas infection rates in percent for 28 missense variants classified as VVCC. Linear correlation was insignificant (Pearson Coefficient $r = 0.13$, Spearman Coefficient $\rho = 0.28$).

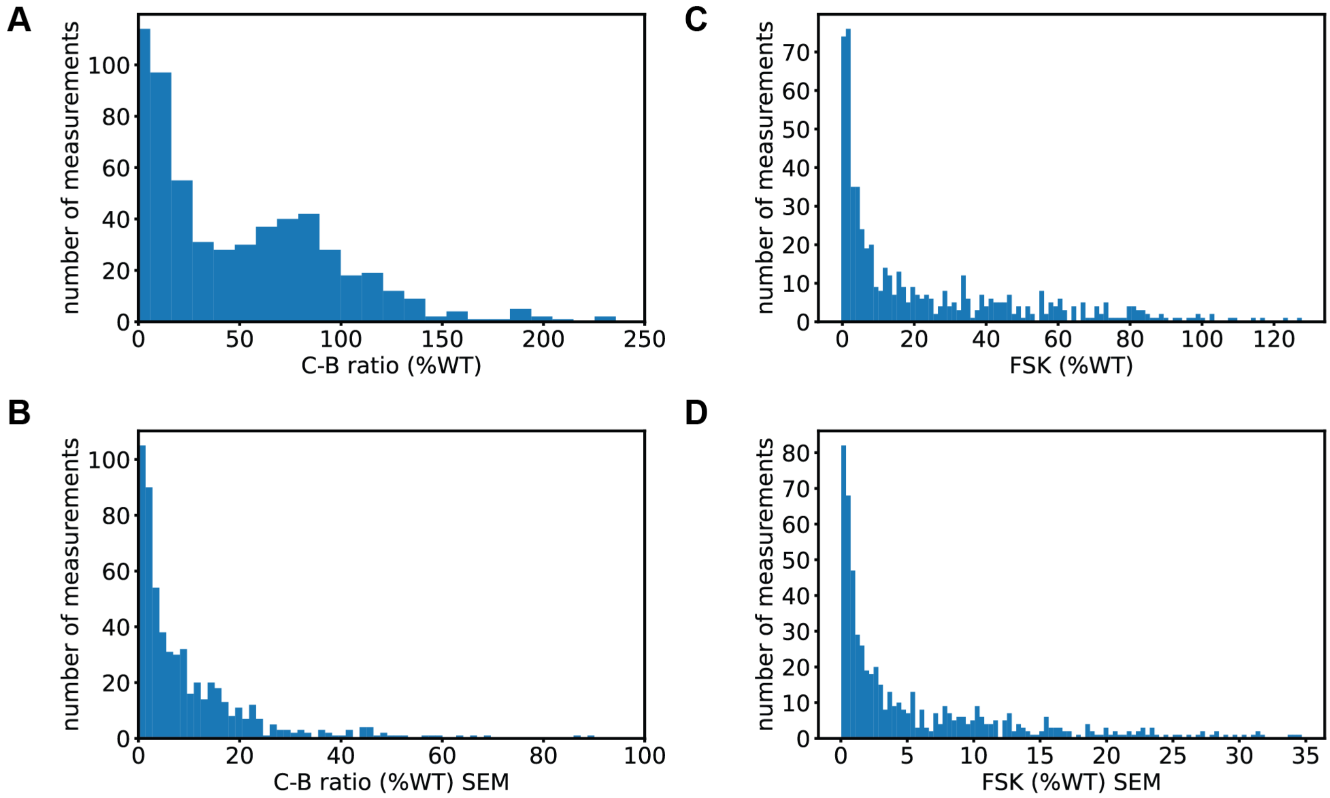


**Supplemental Figure S4. Distributions of experimental data and error from Bihler et al. study for filtering purposes**

**A.** Histogram of the distribution of C-B band ratio of all 585 missense variants from the Bihler et al. study[10].

**B.** Histogram of the distribution of C-B band ratio SEM for all 585 missense variants from the Bihler et al. study. Variants with an SEM greater than 30 were excluded from analysis due to lack of experimental reproducibility and for plotting clarity.

**C.** FSK %WT distribution plotted as a histogram for all 585 missense variants from the Bihler et al. study[10].

**D.** FSK %WT SEM distribution plotted as a histogram for all 585 missense variants from the Bihler et al. study. Variants with an SEM greater than 20 were excluded from analysis due to lack of experimental reproducibility and for plotting clarity.

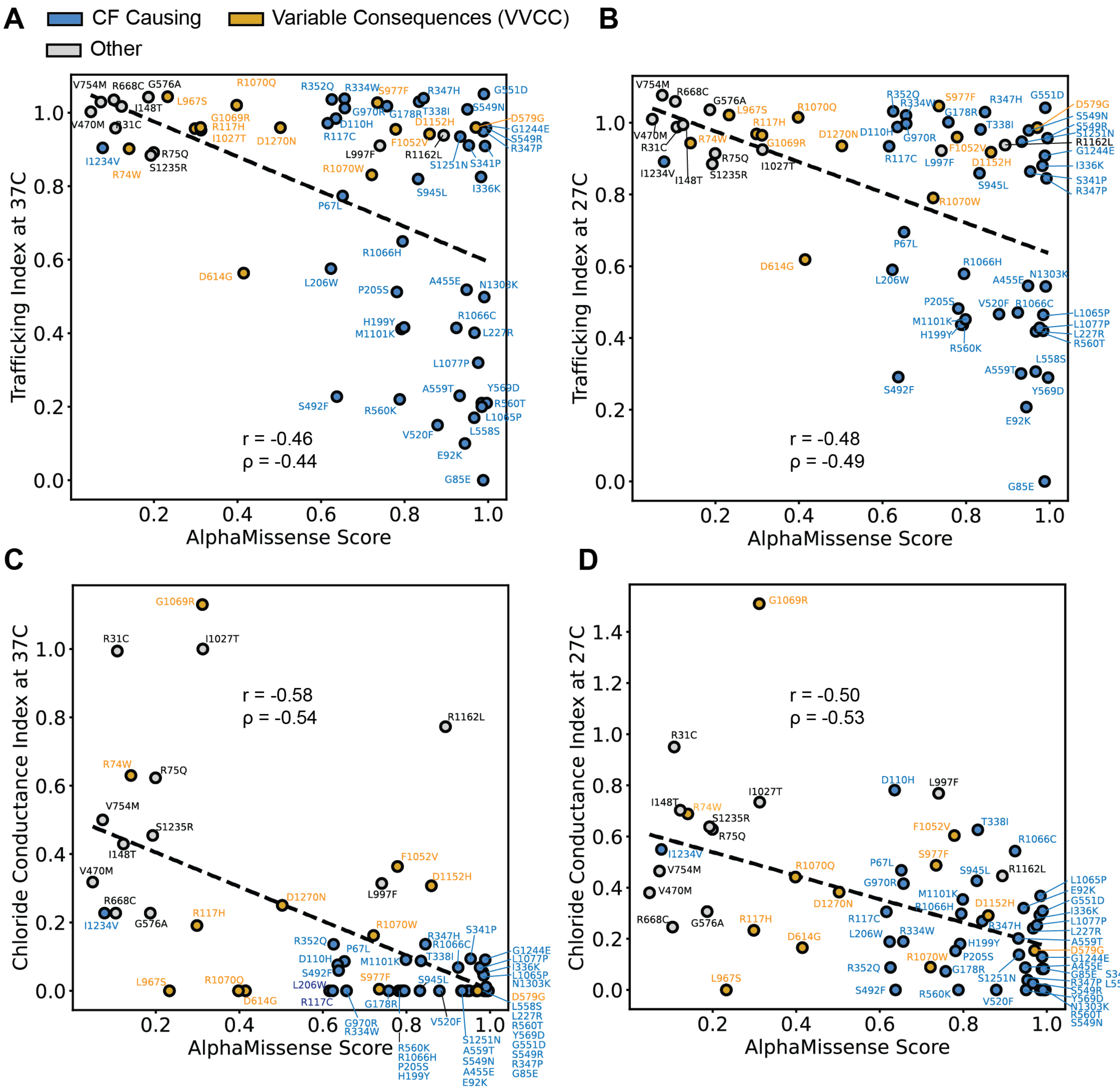

**Supplemental Figure S5. AlphaMissense correlation with CFTR *in vitro* data from spatial covariance study.**
**A.** Spatial covariance data from a previous study[11] for 62 missense variants plotted against AlphaMissense scores. Y axis represents the trafficking index as measured by a western blot trafficking assay when HEK293T cells were incubated at 37 ºC. A slight inverse linear correlation was observed (Pearson Coefficient r = -0.46, Spearman Coefficient ρ = -0.44).
**B.** Spatial covariance data for 62 missense variants using the same trafficking index in **A.** except at 27 ºC, plotted against AlphaMissense scores. Again, an inverse linear correlation was observed (Pearson Coefficient r = -0.48, Spearman Coefficient ρ = -0.49) albeit slightly higher than at 37 ºC.
**C.** AlphaMissense scores correlated with the spatial covariance data but using chloride conductance index described in[11], which measured channel activity at 37 ºC. We observed an increased correlation (Pearson Coefficient r = -0.58, Spearman Coefficient ρ = -0.54).

**D.** AlphaMissense scores correlated with chloride conductance index at 27 ºC. We observed a slight correlation (Pearson Coefficient r = -0.50, Spearman Coefficient ρ = -0.53).

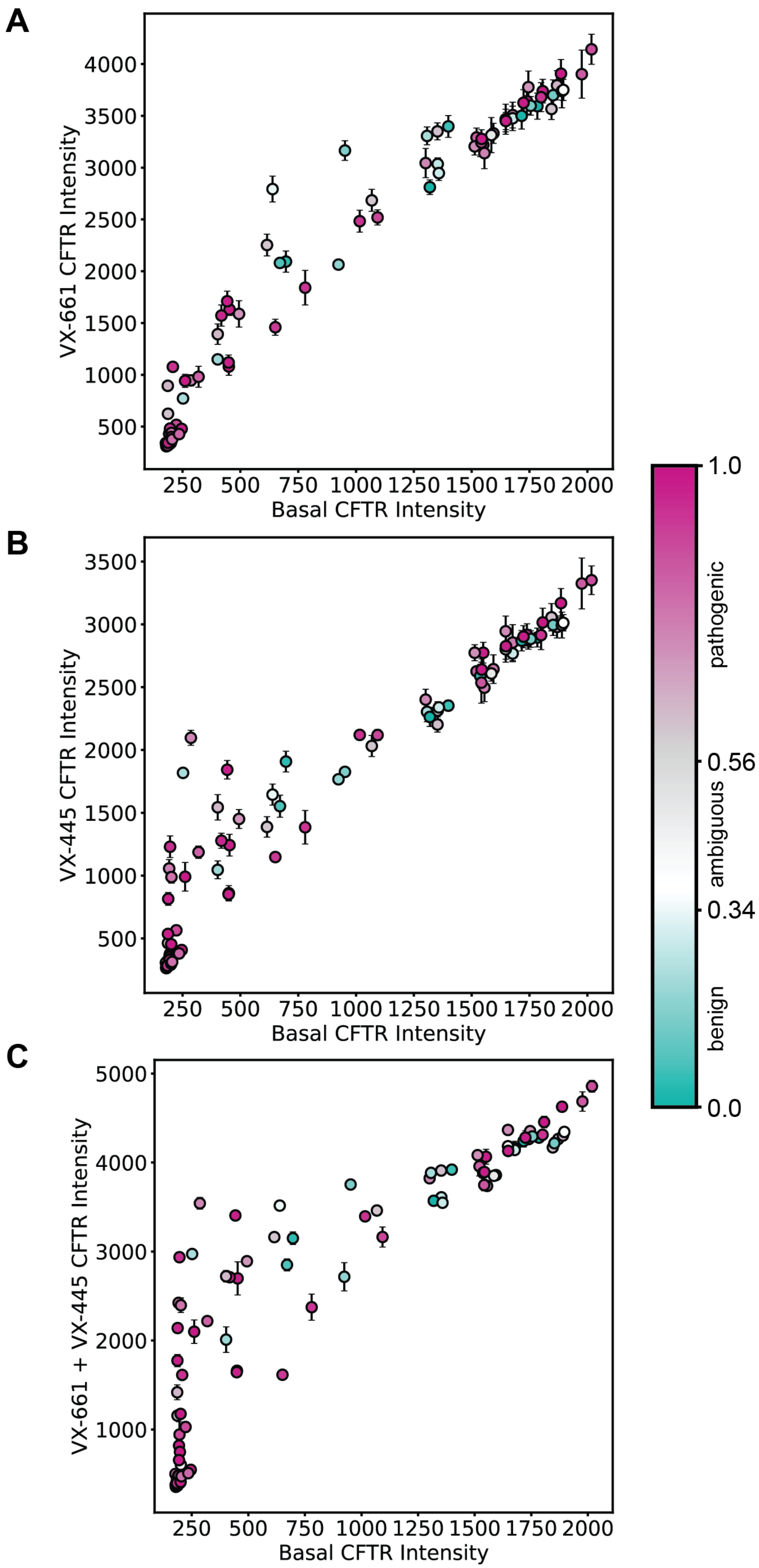
A
VX-661 CFTR Intensity
Basal CFTR Intensity
B
VX-445 CFTR Intensity
Basal CFTR Intensity
C
VX-661 + VX-445 CFTR Intensity
Basal CFTR Intensity
pathogenic
ambiguous
benign
1.0
0.56
0.34
0.0

**Supplemental Figure S6. Deep mutational scanning data for VX-661 and VX-445 response colored by AlphaMissense pathogenicity score.**

**A.** Basal CFTR surface immune staining versus VX-661 CFTR cell surface immune staining intensity[9]. Pathogenic variants score from 0.56-1.00 (violet), ambiguous variants score from 0.34 - 0.56 (grey), and benign variants score 0.04 – 0.34 (green). Error bars represent standard deviation. The distribution of pathogenicity colors throughout the plots suggested that AM pathogenicity prediction score failed to predict the VX-661 response.

**B.** Basal CFTR surface immune staining versus VX-445 CFTR cell surface immune staining intensity. Colored the same as in **A.** Error bars represent standard deviation. Again, AM score failed to predict VX-445 response.

**C.** Basal CFTR surface immune staining versus VX-661 + VX-445 CFTR cell surface immune staining intensity. Colored the same as in **A.** Error bars represent standard deviation. Finally, AM score failed to predict the combination of VX-661 and VX-445 response on a variant basis.

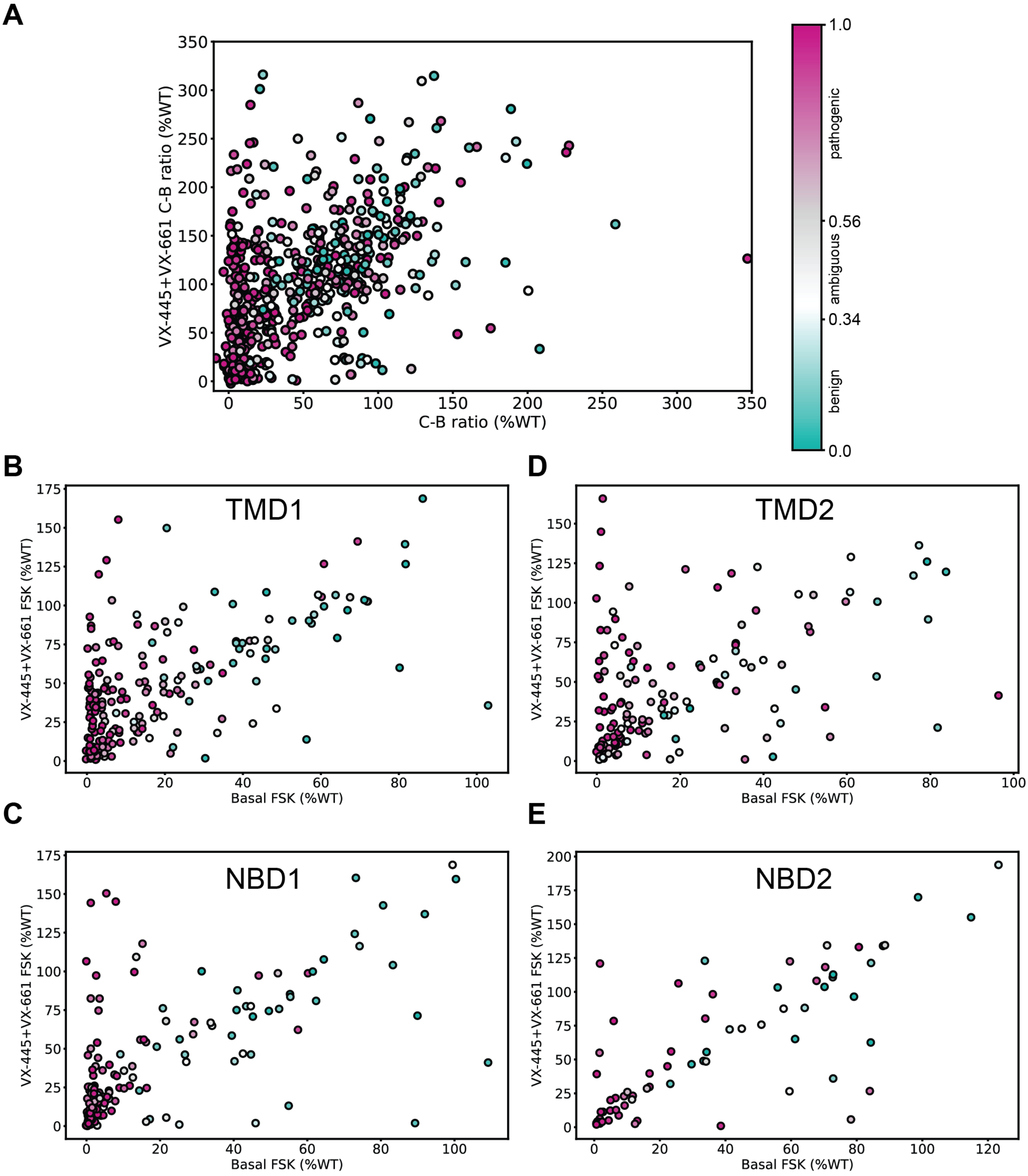


**Supplemental Figure S7. CFTR modulator response plots colored by AlphaMissense pathogenicity score reveals little predictive capabilities of AM in theratyping.**

**A.** Basal mature CFTR (C band) to immature CFTR (B band) trafficking (C-B ratio) in percent WT versus modulator enhanced C/B ratio in percent WT from the Bihler et al. study[10]. Error bars were excluded for clarity. Variants with an AlphaMissense pathogenicity prediction score from 0.56-1.00 were classified by AM as

pathogenic (violet), a score from 0.34 - 0.56 as ambiguous (grey), and a score from 0.04 – 0.34 as benign (green). The distribution of colors across the plots indicated little predictive capability of AM on trafficking theratype.
**B.** TMD1 variants only from **Figure 4B.** of the basal FSK CFTR activity in percent WT versus modulator enhanced FSK CFTR activity in percent WT from the Bihler et al. study[10]. Error bars were excluded for clarity and AM predicted pathogenicity colored as in **A.**
**C.** NBD1 variants only from **Figure 4B.**, data from the Bihler et al. study[10].
**D.** TMD2 variants only from **Figure 4B.**, data from the Bihler et al. study[10].
**E.** NBD2 variants only from **Figure 4B.**, data from the Bihler et al. study[10].
